# Distinct roles of core autophagy-related genes (ATGs) in zebrafish definitive hematopoiesis

**DOI:** 10.1101/2022.01.03.474766

**Authors:** Xiang-Ke Chen, Zhen-Ni Yi, Jack Jark-Yin Lau, Alvin Chun-Hang Ma

**Author notes:** **Address for Correspondence:** Dr. Alvin Chun-Hang MA, PhD; Department of Health Technology and Informatics, Rm. Y924, Lee Shau Kee Building, The Hong Kong Polytechnic University, Hung Hom, Hong Kong.

## Abstract

Despite the well-described discrepancy between some of the macroautophagy/autophagy-related genes (ATGs) in the regulation of hematopoiesis, the varying essentiality of core ATGs in vertebrate definitive hematopoiesis remains largely unclear. Here, we employed zebrafish (*Danio rerio*) to compare the function of six core *atgs* from the core autophagy machineries, which included *atg13, beclin1* (*becn1*), *atg9a, atg2a, atg5*, and *atg3*, in vertebrate definitive hematopoiesis via CRISPR-Cas9 ribonucleoprotein targeting. Zebrafish embryos with various *atg* mutations showed autophagic deficiency throughout the body, including hematopoietic cells. The *atgs* mutations unsurprisingly caused distinctive hematopoietic abnormalities in zebrafish. Notably, *becn1* or *atg9a* mutation resulted in hematopoietic stem cells (HSCs) expansion during the development of the embryo into a larva, which can be attributed to the proteomic changes in metabolism, HSCs regulators, and apoptosis. Besides, *atg3* mutation lowered the leukocytes in developing zebrafish embryos. Intriguingly, a synergistic effect on HSCs expansion was identified in *atg13+becn1* and *atg9a+atg2a* or *atg3* double mutations, in which *atg13* mutation and *atg2a* or *atg3* mutation exacerbated and mitigated the HSCs expansion in *becn1* and *atg9a* mutations, respectively. In addition, the myeloid cell type-specific effects of various *atgs* were also determined between neutrophils and macrophages. Of these, a skewed ratio of neutrophils versus macrophages was found in *atg13* mutation, while both of them were reduced in *atg3* mutation. These findings demonstrated the distinct roles of *atgs* and their interplays in zebrafish definitive hematopoiesis, thereby suggested that the vertebrate definitive hematopoiesis is regulated in an *atgs*-dependent manner.

## Introduction

Macroautophagy (hereinafter autophagy) serves as the scavenger of the cells by removal of harmful components via the lysosomal degradation, which, as an essential but complex process, is tightly regulated by a category of genes, namely autophagy-related genes (ATGs) [1]. Approximately 20 core ATGs that orchestrate the critical steps of canonical autophagy were classified into six functional groups/ machineries, including the unc-51 like autophagy activating kinase (ULK) 1 complex (initiation), class 3 phosphoinositide 3-kinase (PI3K) complex (isolation membrane nucleation), ATG9a vesicles (providing membrane), ATG2a complex (membrane elongation), ATG12 conjugation system (membrane elongation and linking microtubule-associated protein 1A/1B-light chain 3 [LC3] with PE), and LC3-PE conjugation system (membrane elongation and target recognition), which are highly conserved across the eukaryotes [2]. Loss of core ATGs commonly results in neonatal lethality in mice, while the exact cause of death remains elusive [3]. Despite the well-known essentiality of core ATGs in mice, a recent clinical study reported the identification of twelve patients from five families who survived with severely impaired autophagy due to the loss of ATG7, one of the most well studied core ATGs [4]. This was ascribed to the non-canonical or alternative autophagy that undergoes in the absence of some core ATGs since the autophagosome could still be formed [4,5]. Many core ATGs-independent autophagy, such as those that bypass ULK1, Beclin1 (BECN1), ATG5, and ATG7, as well as the autophagy-independent functions of core ATGs have been identified in the past decade [6,7]. Nevertheless, the majority of previous studies still focused on canonical autophagy, and the distinctive effects of various core ATGs are largely neglected.

Previous studies have shown the resemblance and discrepancy between core ATGs in the regulation of hematopoiesis, which is a vital biological process of blood cellular components formation, although very few of them included more than one ATG [8]. In definitive hematopoiesis, ATG5, ATG7, or FAK family-interacting protein of 200 kDa (FIP200) conditional knockout (CKO) in hematopoietic cells declined the number of hematopoietic stem cells (HSCs) [9–11], whereas BECN1 and ATG12 CKO failed to alter the total number of HSCs [12,13]. Moreover, ATG5 or ATG7 CKO reduced the number of multi-lineage progenitors [9,10]; in contrast, their number increased and remained unchanged in ATG12 and BECN1 CKO, respectively [12,13]. Furthermore, ATG7, ATG12, or FIP200 CKO but not BECN1 as well as ATG5 CKO resulted in myeloproliferation, while ATG5, ATG7, or FIP200 CKO but not ATG12 or BECN1 CKO caused anemia [9–13]. These differential hematopoietic abnormalities found in these CKO of the core ATGs demonstrated the core ATGs-dependency of hematopoiesis, while more studies on other core ATGs are needed. In addition to the differences between ATGs, contradictory results were also observed in core ATGs CKO, including ATG5 and ATG7 [9,13,14], which indicate the important ‘timing’ of various core ATGs in the regulation of hematopoiesis. For instance, ATG7 is indispensable for adult but not neonatal HSCs in mice [14]. However, whether this ‘timing’ also occurs in other core ATGs or not remains unexplored.

Over the past decades, zebrafish (*Danio rerio*), a tropical freshwater fish, has emerged as a vital genetic model in the field of hematopoiesis as well as autophagy due to its genetic tractability, small and transparent body, *in vitro* embryogenesis, and, more importantly, evolutionarily conversed genes orthologous to 70% of the human genes [15–17]. With the advanced gene editing tools, such as Clustered Regularly Interspaced Short Palindromic Repeats (CRISPR)-Cas9 ribonucleoprotein (RNP) complex [18], zebrafish embryos has gradually become a more efficient, convenient, and feasible model than mice in genetic screening *in vivo*, particularly in the study of hematopoiesis. Nevertheless, the roles of core *atgs* in zebrafish hematopoiesis remain largely unknown. Here, we reported for the first time, the differential effects of *atgs* mutations via CRISPR-Cas9 RNP targeting on zebrafish definitive hematopoiesis, although their mutations all disturbed the autophagy process, which highlighted the essentiality of various *atgs*-dependent effects rather than the uniformed canonical autophagy-dependent effect in vertebrate definitive hematopoiesis.

## Results

### *Core* atgs *targeting by CRISPR-Cas9 ribonucleoprotein (RNP)*

Core ATGs that are required for autophagy process have been classified in to six evolutionarily conserved functional groups, also known as ‘core autophagy machineries’ [2]. In this study, core *atgs* was selected form the six core machineries of autophagy for CRISPR-Cas9 sgRNP complex targeting, including *atg13* (ULK1 complex), *becn1* (PI3K complex), *atg9a* (ATG9a vesicles), *atg2a* (atg2a complex), *atg5* (ATG12 conjugation system), and *atg3* (LC3-PE conjugation system) (**Figure 1A-B**). Phylogenetic analysis revealed that these *atgs* were conserved between zebrafish and humans with high similarities (**Figure S1A-D**). crRNA were designed for each *atg* targeting early exon to model null-like mutation in zebrafish embryos (**Figure 1C**). Very high mutagenesis efficiencies (> 95%) were observed in all somatic *atgs* mutants as shown by restriction fragment length polymorphism (RFLP) assay (**Figure 1F**) and Sanger sequencing (**Figure S1E**), which can result in a null-like loss of *atg* protein (**Figure S2A**). In addition, a stable production of null-like zebrafish mutants can be achieved by the skilled operator (**Figure S2B**). While most of the mutant embryos (> 75%) displayed normal development and morphology in embryonic stages **(Figure 1D**), deformed embryos could also been found (**Figure 1E**). However, most of these mutants, except *atg9a* and *atg2a* mutants can only survive up to around 2 week-post-fertilization.

**Figure 1.**
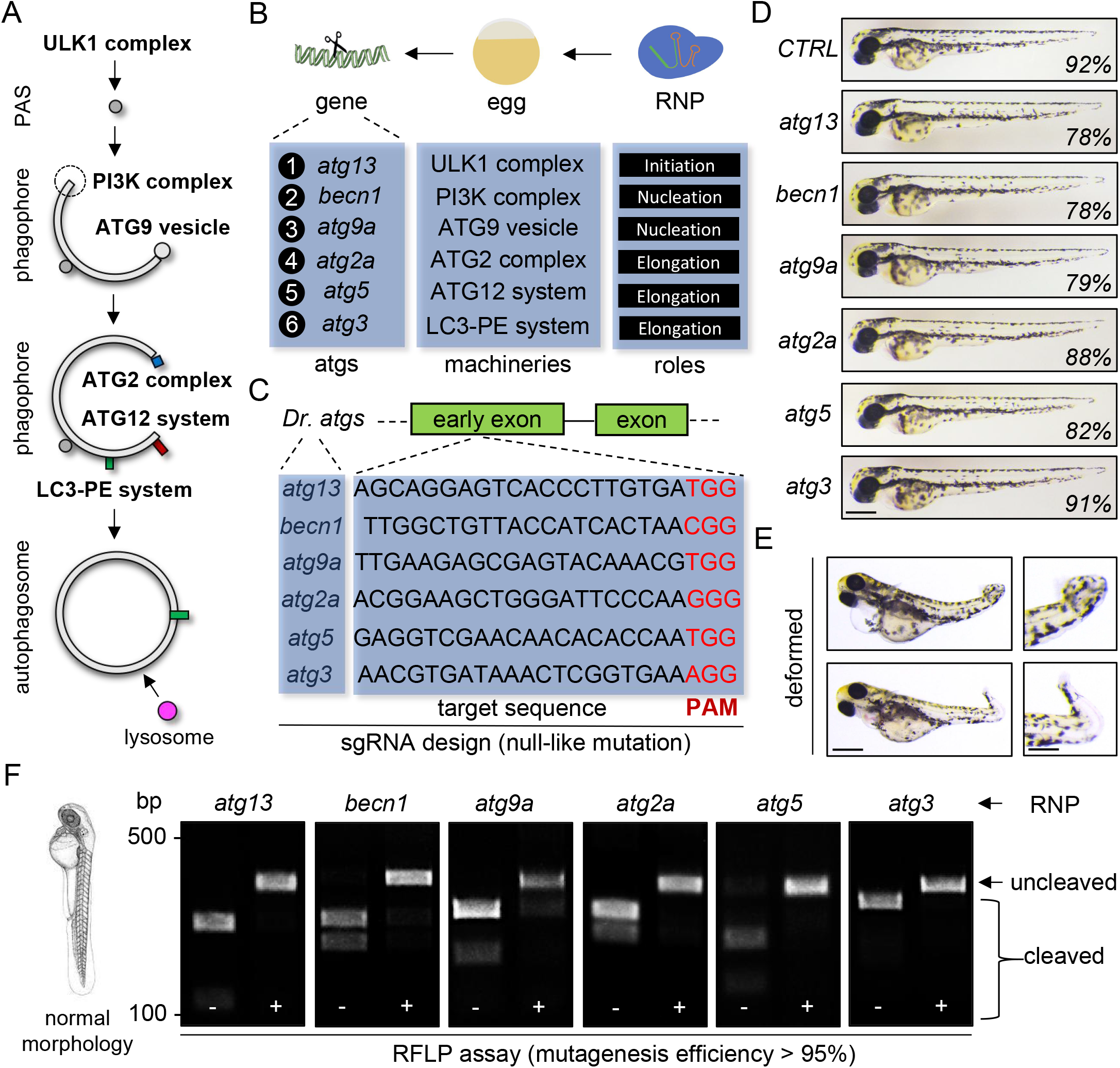
Core atgs targeting by CRISPR-Cas9 ribonucleoprotein (RNP). (**A**) Schematic diagram showing the involvement of autophagy machineries in autophagy pathway. PAS, pre-autophagosomal structure. (**B**) Schematic diagram showing the CRISPR-Cas9 RNP targeting and core autophagy-related genes (*atgs*) selected from autophagy machinery (cam). (**C**) Target sequences and sgRNA design of various *atgs*. (**D**) Morphology and percentage of normal morphology of zebrafish with *atgs* mutations. Scale bar, 0.5 mm. (**E**) Representative deformed zebrafish embryos with *atg* mutants. Scale bar, 0.5 mm. (**F**) Restriction fragment length polymorphism (RFLP) assay and mutation efficiency.

### *Autophagy deficiency in core* atgs *mutant zebrafish embryos*

We next examined autophagy after *atgs* targeting in Tg(Lc3:GFP) zebrafish line. The number of Lc3+ autophagy puncta (autophagosome) in the muscle was significantly lowered after *atg13, becn1, atg9a, atg5*, and *atg3* targeting but not in *atg2a* targeting (**Figure 2A-C**). The inconsistent autophagic activation or basal level was not only detected among various *atgs* mutations but also among various organs and tissues (**Figure S2C**). Intriguingly, all the *atgs* mutations cannot completely block autophagy, which suggested the presence of individual *atg*-independent alternative autophagy. In addition, autophagy flux was measured upon Chloroquine (CQ) treatment. An elevated number of puncta was observed in the control (CTRL), *becn1, atg9a*, and *atg5* mutants after CQ treatment, while it remained unchanged in *atg13, atg2a*, and *atg3* mutants (**Figure 2A-C**). In particular, the number of puncta in *atg2a* mutants remain unchanged with or without CQ treatment, which suggested that its primary function in the blockage of autophagic flux. However, its mechanism is distinct from CQ, where the lysosome can still fuse with autophagosome but cannot undergo subsequent degradation (**Figure 2D-G**). Inconsistent with the observed autophagic changes in muscle cells, western blot of whole embryos showed a similar reduction in the *Lc3II* level in various *atgs* mutants other than *atg13*, which also indicated the tissue-specific autophagic changes (**Figure 2H-I**). A more reliable and sensitive CytoID staining [19] was performed in the sorted *coro1a+* hematopoietic cells since very few puncta and Lc3 signal can be detected in the *coro1a+* cells of Tg(Lc3:GFP;coro1a:DsRed) zebrafish line (**Figure S2D**) and CytoID staining failed to penetrate the live embryo. Importantly, CytoID+ autophagic vacuoles declined in almost all *atgs* mutants similar to the decreased number of puncta in the muscle (**Figure 3A-C**), demonstrated that *atgs* mutations resulted in autophagy deficiency in the hematopoietic cells.

**Figure 2.**
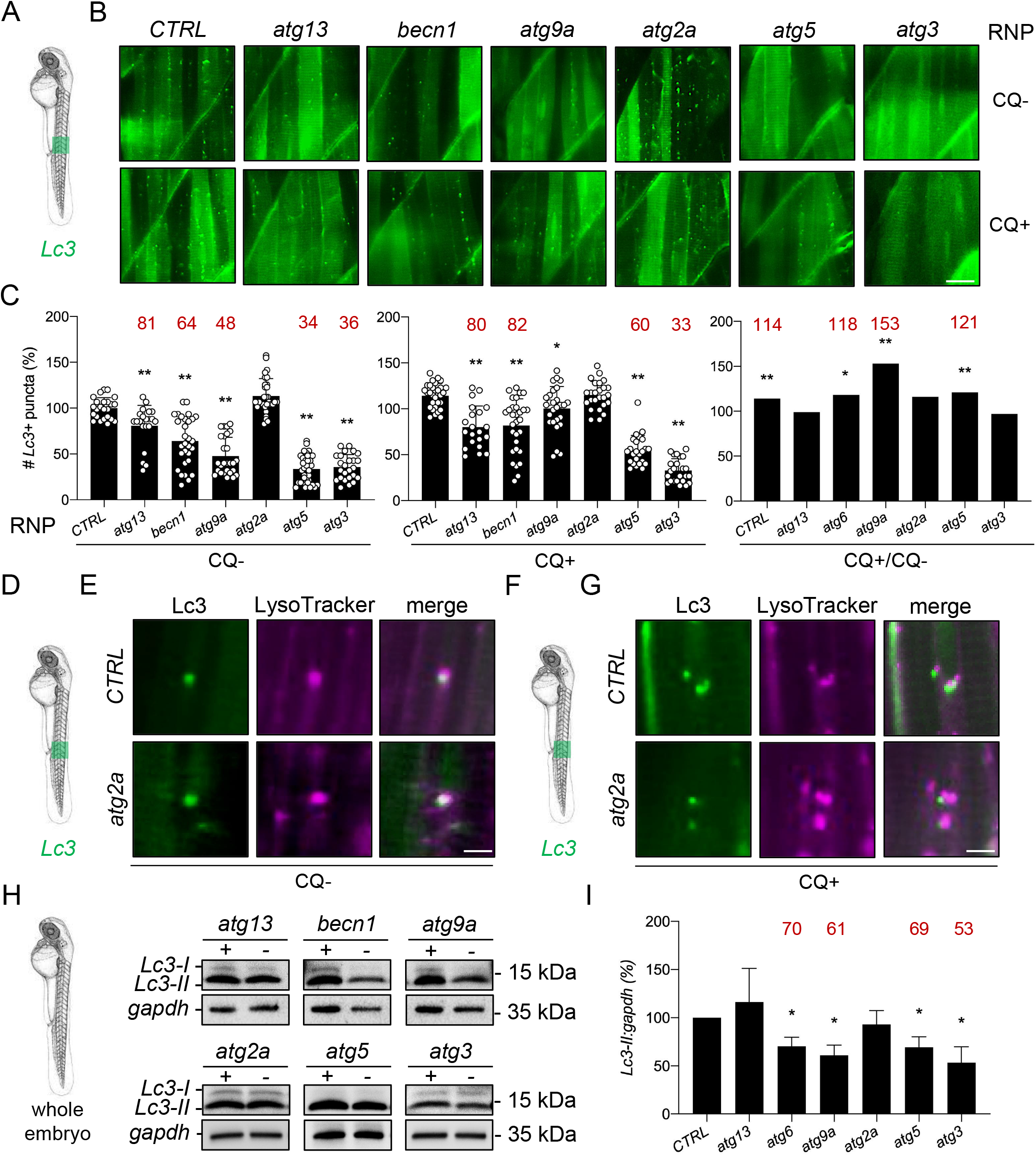
Autophagic deficiency in zebrafish embryos with core *atgs* mutations. (**A-C**) Autophagosomes or Lc3+ puncta in the muscle of Tg(Lc3:GFP) zebrafish embryos with (+) and without (-) Chloroquine (CQ) treatment in various *atgs* mutation. *, p < 0.05 compared with CTRL or CQ-. **, p < 0.01 compared with CTRL or CQ-. Scale bar, 50 μm. (**D-G**) Fusion of autolysosome between Lc3+ autophagosome and LysoTracker+ lysosome in Tg(Lc3:GFP) zebrafish embryos with (+) and without (-) CQ treatment in various *atgs* mutation. Scale bar, 5 μm. (**H-I**) Western blotting result of *Lc3* and *gapdh* protein levels in zebrafish embryos with various *atgs* mutation.*, p < 0.05 compared with CTRL.

**Figure 3.**
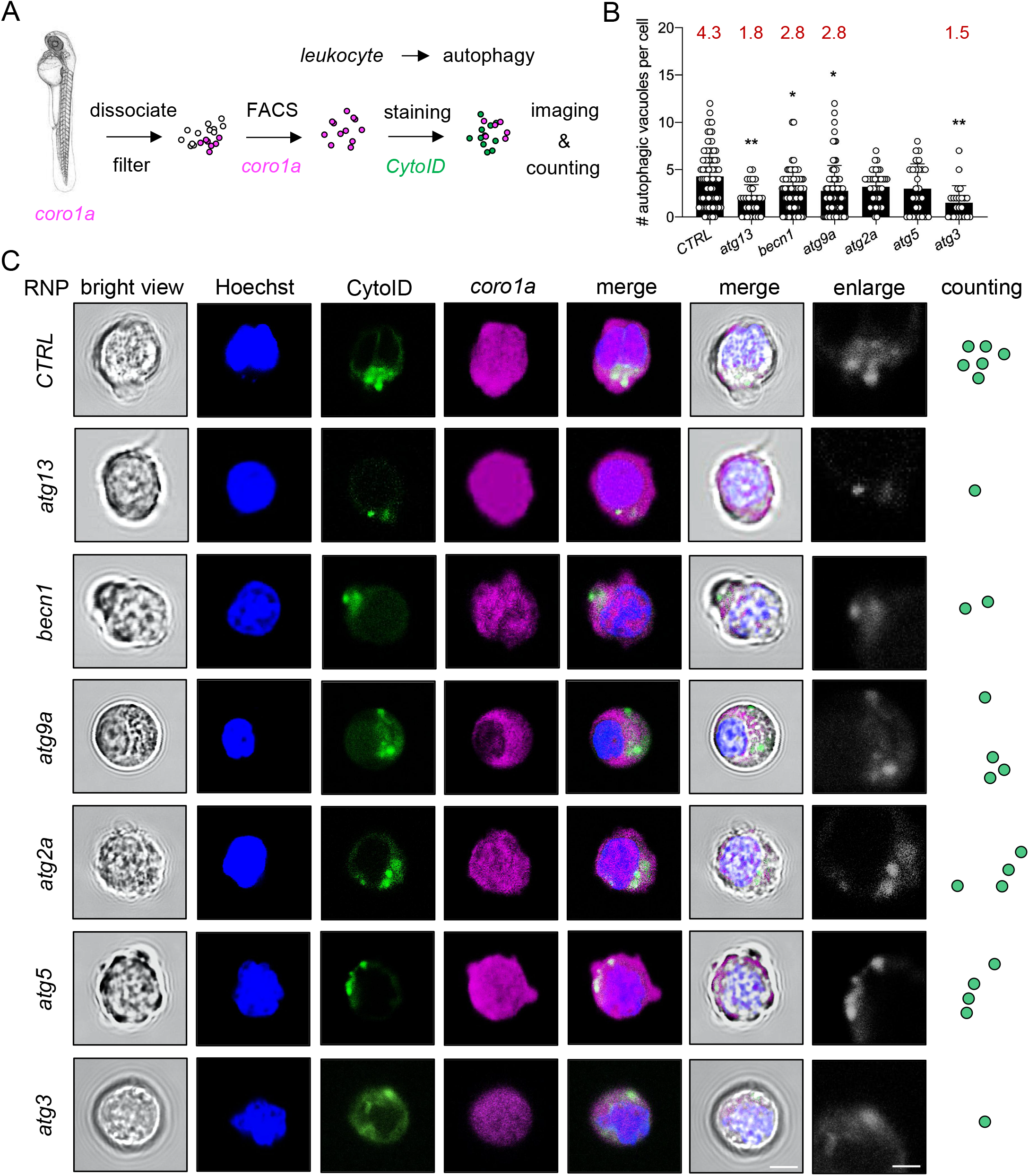
Attenuation of autophagic vacuoles in leukocytes with core *atgs* mutations. (**A**) Experimental setup for Cyto-ID+ autophagic vacuoles measurement in *coro1a+* leukocytes. (**B-C**) Representative images and quantification of Cyto-ID+ autophagic vacuoles in *coro1a+* leukocytes sorted from zebrafish embryos with various *atgs* mutation. *, p < 0.05 compared with CTRL. **, p < 0.01 compared with CTRL. Scale bar, 5 μm.

### *Distinct effects of* atgs *mutation on definitive hematopoiesis*

We then examined the effects of *atgs* mutations on definitive hematopoiesis by whole-mount in suit hybridization (WISH). While all the *atgs* mutations resulted in autophagy deficiency, only *atg13, becn1*, and *atg3* mutations affected the number of *c-myb+* HSCs and *lcp1+* pan-leukocytes in the caudal hematopoietic tissue (CHT), albeit modestly (< 30% of hematopoietic cells) (**Figure 4A-C and 4G-I**). More importantly, *atg13* and *atg3* mutations reduced but *becn1* mutation increased *c-myb+* HSCs, while *atg13* and *becn1* increased but *atg3* decreased the *lcp1+* pan-leukocytes. These differential hematopoietic phenotypes supported that the effects of *atgs* mutations on HSCs and leukocytes are possibly canonical autophagy-independent. In contrast, *spi1+* myeloid progenitor declined in all the *atgs* mutants probably through the regulation of canonical autophagy, though more evidence is needed (**Figure 4D-F**). In addition, no significant difference was observed in the *hbae1+* erythrocytes in the CHT, which also suggested the hematopoietic linage-specific effect of *atgs* (**Figure 4J-L**). To elucidate the differential effects of *atgs* mutation in definitive hematopoiesis, mass spectrometry-based proteomic analysis was performed to identify the proteomic differences underlying the inconsistent phenotypes (**Figure 5A**). Strikingly, key proteins identified in proteomic analysis (p < 0.05, fold change > 1.50 or < 0.66) varied between *atgs* mutations, in which *beclin1* mutations have the largest number of altered proteins comparing with CTRL (**Figure 5B-C**).

**Figure 4.**
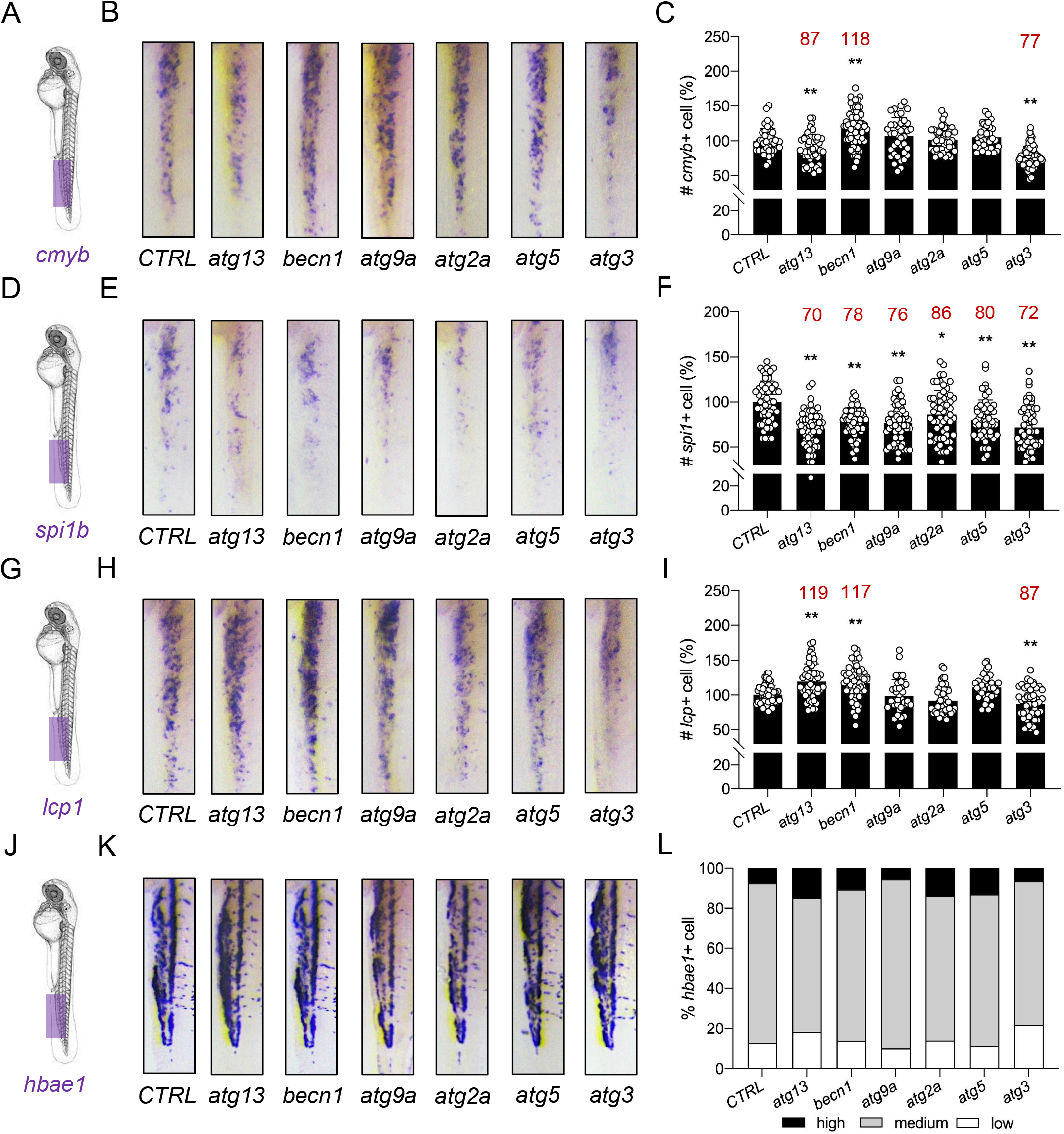
Distinct effects of core *atgs* mutation on definitive hematopoiesis in zebrafish. (**A-C**) Whole mount in situ hybridization (WISH) results of *cmyb+* HSCs. **, p < 0.01 compared with CTRL. (**D-F**) WISH results of *spi1b+* myeloid progenitor. *, p < 0.05 compared with CTRL. **, p < 0.01 compared with CTRL. (**G-I**) WISH results of *lcp1+* pan-leukocytes. **, p < 0.01 compared with CTRL. (**J-K**) WISH results of *hbae1+* erythrocytes. No significant difference was found between CTRL and *atgs* mutations (*C^2^* test). All caudal hematopoietic tissue (CHT) from WISH pictures was straightened by ImageJ.

**Figure 5.**
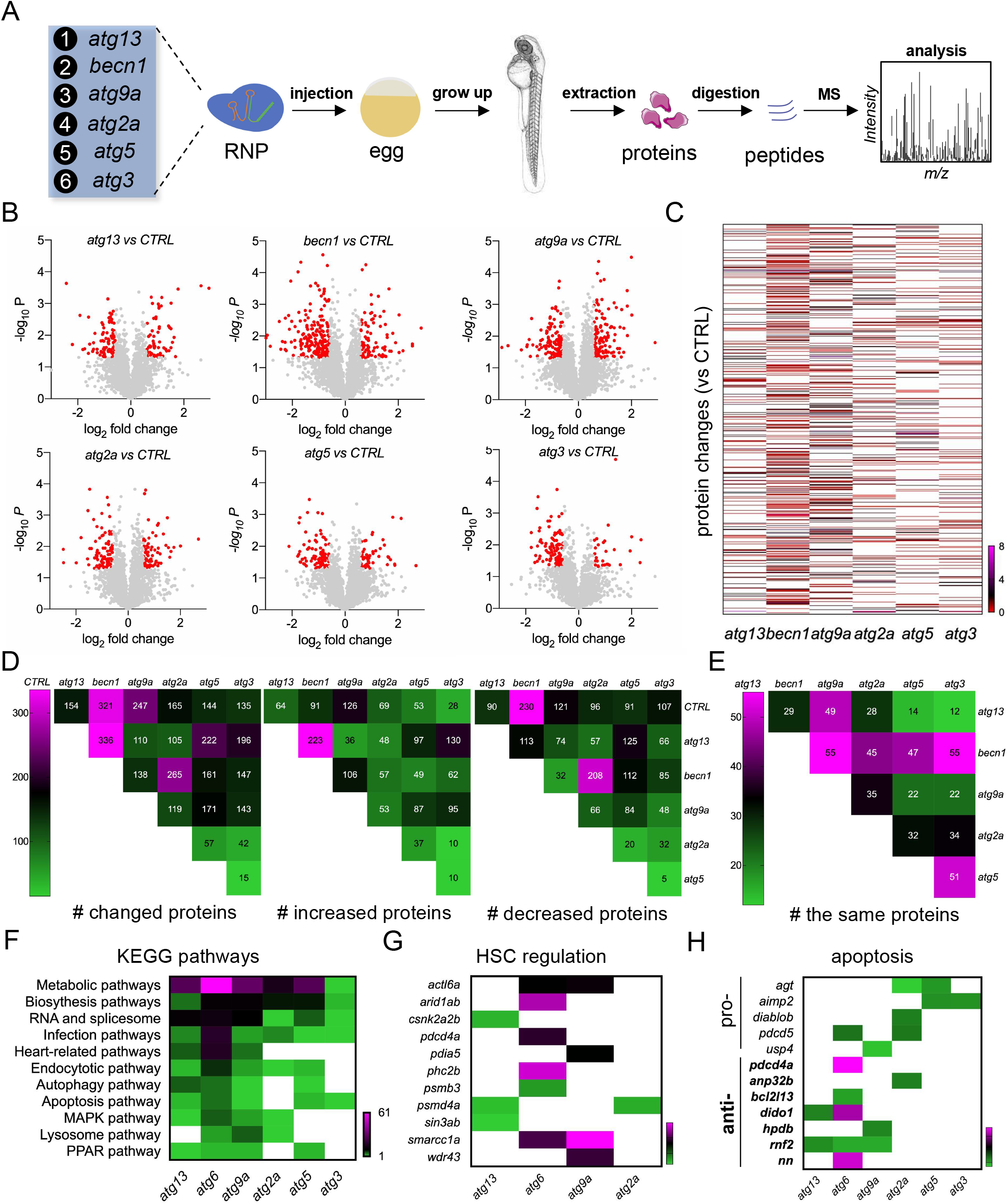
Mass spectrometry-based proteomic variability among *atgs* mutations. (**A**) Experimental setup for mass spectrometry-based proteomic analysis of zebrafish embryos with various *atgs* mutations. (**B-C**) Volcanic map and heat map of comparison between *atgs* mutation and CTRL. Red dots, p < 0.05 and fold change > 1.5. (**D-E**) Comparison of the number (#) of changed proteins, increased proteins, decreased proteins, and the same protein among *atgs* mutation. (**F-H**) Comparison of KEGG pathways and specific proteins (HSCs and apoptosis-related) among *atgs* mutation.

We further compared the proteomic profiles from different *atgs* mutants (**Figure 5D-E and Figure S3**). Interestingly, relatively more proteomic changes were observed in *atg13, beclin1*, and *atg5* mutants, while relatively less proteomic changes were found among *atg2a, atg5*, and *atg3* mutants. KEGG pathways enrichment suggested that metabolic pathways, biosynthesis pathways, and RNA and splicesome are the top three pathways that were changed in *atgs* mutations (**Figure 5F**). Among the *atgs*, less pathways were found in *atg3, atg2a*, and *atg5* mutants compared to *becn1, atg9a*, and *atg13*. These pathways are potentially cooperated to influence definitive hematopoiesis under *atgs* mutants and autophagy deficiency. In particular, a number of HSC regulation-related proteins previously reported were also detected in the proteomic analysis, which were predominantly changed in *becn1* and *atg9a* mutants, while remained mostly unchanged in *atg5* and *atg3* mutants (**Figure 5G**). Similarly, differential proteomic patterns can also be found in apoptosis-related proteins, which indicated an inhibition of apoptosis in zebrafish embryos under *atgs* mutations, especially under *becn1* mutation (**Figure 5H**). Therefore, the roles of *atgs* in definitive hematopoiesis could be, at least partially, attributed to specific proteins that modulate the programmed death, cell cycle, proliferation, and differentiation of hematopoiesis cells, which warrants further investigations.

### *Time-dependent effects of* atgs *double mutations*

To further delineate the effects of *atgs* mutations on definitive hematopoiesis, we co-mutated core *atgs* selectively based on their differential proteomic profiles by co-injecting two sgRNPs targeting two *atgs* and high mutagenic efficiencies similar to single sgRNP injection were obtained (**Figure 6A and Figure S2E**). Since *atgs* has been found to be activated rhythmically in a clock-dependent manner, featuring the dynamic mRNA level of *atgs* during the days after birth [20], we also tracked the effect of *atgs* mutations on definitive hematopoiesis from 2 to 4 dpf to examine the time-dependent effects of *atgs* in definitive hematopoiesis.

**Figure 6.**
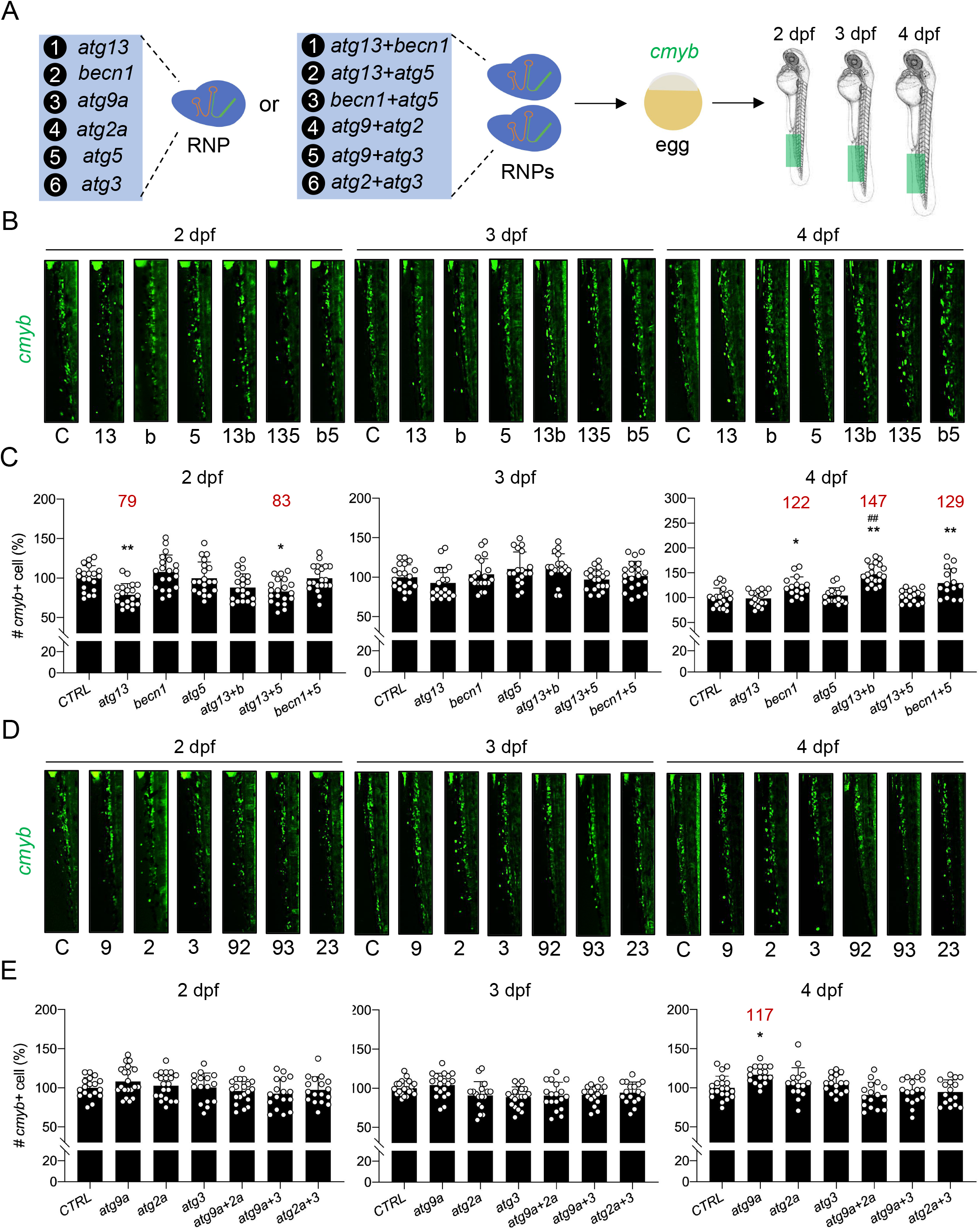
Time-dependent effects of core *atgs* single or double mutations on HSCs. (**A**) Experimental setup for the time-dependent responses of *cmyb+* HSCs to single or double mutations of *atgs*. (**B-C**) Effect of combination among *atg13, becn1*, and *atg5* mutations on *cmyb+* HSCs during the period from 2 dpf to 4 dpf. *, p < 0.05 compared with CTRL. **, p < 0.01 compared with CTRL. ##, p < 0.01 compared with *becn1* mutation. (**D-E**). Effect of combination among *atg9a, atg2a*, and *atg3* mutations on *cmyb+* HSCs during the period from 2 dpf to 4 dpf. *, p < 0.05 compared with CTRL.

Notably, *becn1, atg13+becn1*, and *becn1+atg5* mutations resulted in aberrant accumulation of *cmyb+* HSCs at 4 dpf, where a interaction effect was found in *becn1+atg13* (F = 4.27, *p* < 0.05, two-way ANOVA) but not in *becn1+atg5* (*p* > 0.05, two-way ANOVA) (**Figure 6B-C**). Besides, *atg13* and *atg13+atg5* mutations only declined the number of *cmyb+* HSCs at 2 dpf. While *atg9a* mutation also increased the *cmyb+* HSCs at 4 dpf, which can be alleviated by co-introduction of *atg2a* (F = 18.02, *p* < 0.01, two-way ANOVA) or *atg3* (F = 16.25, *p* < 0.01, two-way ANOVA) mutation (**Figure 6D-E**). Other single injections or co-injections showed no effects on *cmyb+* HSCs. For coro1a+ leukocytes, although *atg13, becn1, atg5*, and *atg13+atg5* mutations leaded to increased *coro1a+* leukocytes, *atg13+becn1* showed an opposite effect, in which *atg13+becn1* attenuated the number of *coro1a+* leukocytes (**Figure 7A-C**). These results indicated the different mechanisms underlying the increased *coro1a+* leukocytes in *becn1* and *atg5* or *atg13* mutants. While the increased *coro1a+* leukocytes in *atg13, becn1*, and *atg5* mutants were no longer observed on 4 dpf, both effects of *atg13+becn1* and *atg13+atg5* co-mutation on leukocytes lasted to 4 dpf though interaction effect was only observed in *atg13+becn1* (F = 13.70, *p* < 0.01, two-way ANOVA) but not in *atg13+atg5 (p* > 0.05, two-way ANOVA) (**Figure 7B-C**). In addition, *atg3, atg9a+atg2a, atg9a+atg3*, and *atg2a+atg3* mutations resulted in the decreased *coro1a+* leukocytes from 2 to 4 dpf. Main effect of *atg3* (F = 21.78, *p* < 0.01, two-way ANOVA) and *atg2a* (F = 20.80, *p* < 0.01, two-way ANOVA) were observed in context of co-mutation with *atg9a* and no interaction effect (*p* > 0.05, two-way ANOVA) was found (**Figure 7D-E**). Altogether, these results indicate the distinctive molecular or cellular mechanism underlying different core *atgs*, which orchestrate in the regulation of definitive hematopoiesis.

**Figure 7.**
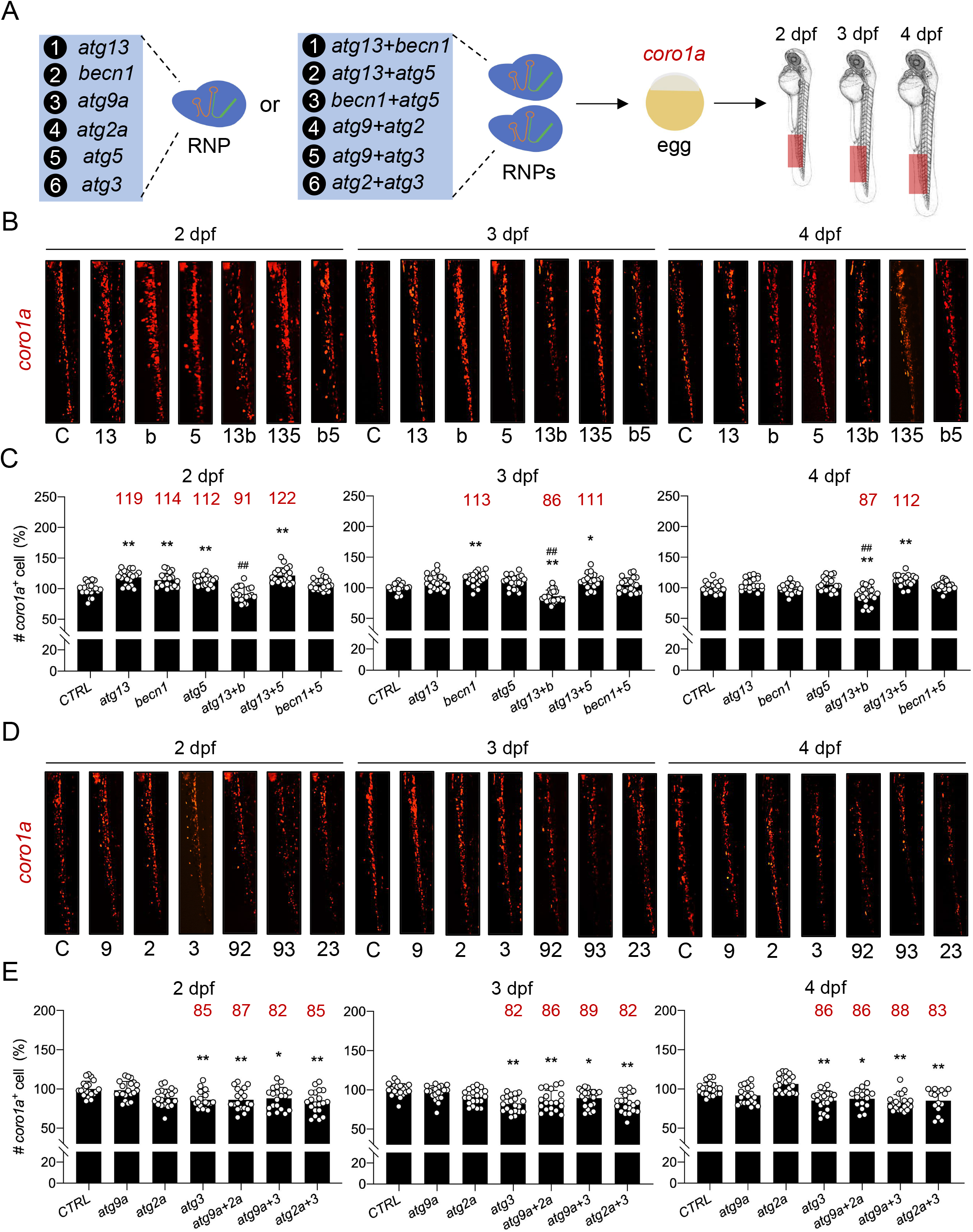
Time-dependent effects of core *atgs* single or double mutations on leukocytes. (**A**) Experimental setup for the time-dependent responses of *coro1a+* leukocytes to single or double mutations of *atgs*. (**B-C**) Effect of combination among *atg13, becn1*, and *atg5* mutations on *coro1a+* leukocytes during the period from 2 dpf to 4 dpf. *, p < 0.05 compared with CTRL. **, p < 0.01 compared with CTRL. ##, p < 0.01 compared with *becn1* mutation. (**D-E**). Effect of combination among *atg9a, atg2a*, and *atg3* mutations on *coro1a+* leukocytes during the period from 2 dpf to 4 dpf. *, p < 0.05 compared with CTRL. **, p < 0.01 compared with CTRL.

### Effects of atgs mutation on myeloid lineages

Comparison of *atgs* expression in various hematopoietic lineages was conducted in both human and zebrafish by *in silico* analysis of published gene expression database, which showed a huge variability and only a certain portion of hematopoietic cells expressed some of the ATGs/*atgs* (**Figure S4A-B**). Therefore, we further examined the effect of core *atgs* mutations on the two major types of myeloid cells, neutrophils and macrophages from 2 to 4 dpf (**Figure 8A**). In accordance with the changes in *coro1a+* leukocytes, *atg13* and *becn1* mutations elevated the number of *mpx+* neutrophils at 2 dpf, while *atg9a, atg2a*, and *atg3* showed the opposite effect (**Figure 8B-C**). In addition, *atg13, atg5*, and *atg3* mutations reduced the number of *mfap4+* macrophages, while no significant difference was found in other *atgs* mutants (**Figure 8D-E**). Interestingly, although the number of *coro1a+* leukocyte was not affected in *atg13* mutant, a shift from *mfap4+* macrophages to *mpx+* neutrophils was observed (**Figure 7B-C and Figure 8A-E**). This indicated the opposite roles of *atg13* in the regulation of neutrophil and macrophage. Conversely, deceased *coro1a+* leukocytes in *atg3* mutant was resulted from both decreased *mpx+* neutrophils and *mfap4+* macrophages (**Figure 7D-E and Figure 8A-E**). Meanwhile, *atgs* mutation that only affected either neutrophil or macrophage population, such as the decreased neutrophils in *atg9a* or *atg2a* mutant and the decreased macrophages in *atg5* mutant, were correlated to the subtle effects on total number of leukocytes (**Figure 7 and Figure 8**). Unlike *cmyb+* HSCs and *coro1a+* leukocytes, almost no time-dependent effect of *atgs* mutations on *mpx+* neutrophils and *mfap4+* macrophages was observed during the development form 2 dpf to 4 dpf in this study.

**Figure 8.**
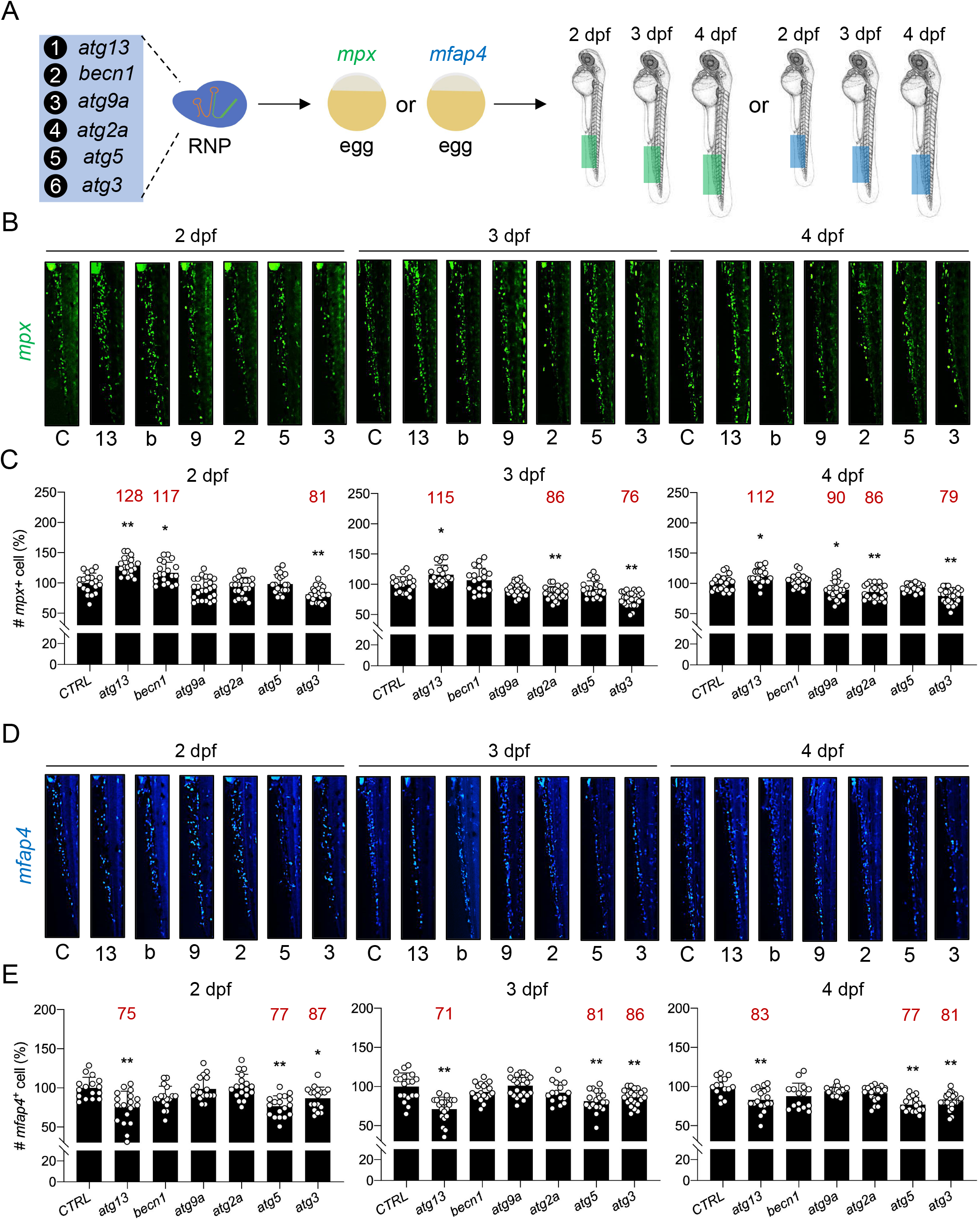
The effect of core *atgs* mutation in myeloid lineages. (**A**) Experimental setup for the time-dependent responses of *mpx+* neutrophils and *mfap4+* macrophages to *atgs* mutation. (**B-C**) Effect of *atg13, becn1, atg9a, atg2a, atg5*, and *atg3* mutations on *mpx+* neutrophils during the period from 2 dpf to 4 dpf. *, p < 0.05 compared with CTRL. **, p < 0.01 compared with CTRL. (**D-E**) Effect of *atg13, becn1, atg9a, atg2a, atg5*, and *atg3* mutations on *mfap4+* macrophages during the period from 2 dpf to 4 dpf. *, p < 0.05 compared with CTRL. **, p < 0.01 compared with CTRL.

## Discussion

Defective canonical autophagy has long been involved in the hematopoietic disturbances observed in core ATGs-deficient mice [21]. The evidence emerging in the past decade, however, reported the distinct effects of various core ATGs ablation on definitive hematopoiesis [9–13]. It suggested a canonical autophagy-independent role of core ATGs, which largely comprise core ATG-dependent non-canonical autophagy and non-autophagy functions, in the hematopoietic system, although very few core ATGs were compared. In this context, we hypothesized that the core ATG has both shared effects with other core ATGs and its own distinctive effects on definitive hematopoiesis. Therefore, we selected six core ATGs from different core autophagy machineries, including *atg13* (*ulk1* complex), *becn1* (PI3K complex), *atg9a* (*atg9a* vesicles), *atg2a* (*atg2a* complex), *atg5* (*atg12* conjugation system), and *atg3* (*Lc3*-PE conjugation system) [2], and then examined their functions in definitive hematopoiesis by using CRISPR-Cas9 RNP targeting in the zebrafish model. We first observed the autophagy deficiency in the body and hematopoietic cells of various *atgs-*mutant zebrafish embryos while it varied among *atgs*. As expected, the autophagic and hematopoietic responses to *atgs* mutations were inconsistent. Mutation of core *atgs* has distinct effects on the definitive hematopoiesis, with some of the effects are time- or myeloid cell type-dependent. Furthermore, the interactions between core *atgs* in definitive hematopoiesis were also determined by double mutations, which revealed a synergistic effect between some of the core *atgs;* and demonstrated the interplays between *atgs* in zebrafish definitive hematopoiesis.

Approximately 20 core ATGs that are fundamental to canonical autophagy have been functionally categorized into six core autophagy machineries [2]. Since the canonical autophagy is essential for the development, mutations of core *atgs* corresponding to various core autophagy machineries except for *atg9a* and *atg2a* in the present study resulted in the larval lethality in accordance with a previous report [22]. Notably, this indicated the similarity of *Atgs/atgs* in the early development between zebrafish and mice, while the exact cause of death needs further investigation [3]. As core ATGs that primarily facilitate the formation of autophagosome, autophagic responses to *atg13, becn1*, and *atg5* knockout (KO) or knockdown (KD) have been previously studied in zebrafish, whereas studies on *atg9a, atg2a*, and *atg3* mutations are not yet reported. Of these, morpholino (MO) KD of *atg13* and *atg5* decreased the level of autophagosome characterized by *Lc3-II* protein level or *Lc3+* puncta in zebrafish [23,24], which are consistent with the reduced autophagosome observed in this study after CRISPR-Cas9 RNP complex-based *atg13* and *atg5* targeting. Conversely, contradictory results were observed in *becn1* mutations, in which MO KD of *becn1*, exon two targeting in this study, and exon four targeting in the other study attenuated the level of autophagosome, while exon seven targeting in another study increased the level of autophagosome in zebrafish [24–26]. These results suggested a mutation type-dependent effect of *becn1* on autophagosome formation, and targeting an early exon of *becn1* may be required for the canonical autophagy deficiency. More importantly, to our knowledge, this is the first study that described autophagy defects in zebrafish embryos with *atg9a, atg2a*, and *atg3* mutations. Like *atg13, becn1*, and *atg5*, the functions of *atg9a, atg2a*, and *atg3* in canonical autophagy of zebrafish are similar to their orthologs in mammalians. Because ATG9a and ATG3 are responsible for autophagosome formation and LC3 lipidation, respectively [27,28], a decreased level of autophagosome was detected in zebrafish with *atg9a* and *atg3* mutations. In addition, loss of ATG2a is associated with impaired autophagic flux and accumulation of immature autophagosomal membranes in mammalian cells [29], which was reproduced in *atg2a*-mutant zebrafish embryos. Besides, *atg13* and *atg3* mutations also affected the autophagic flux in zebrafish through a distinct mechanism by inhibiting the autophagosome formation and LC3 lipidation, respectively [28,30]. Altogether, the function of various *atgs* in the canonical autophagy process conservatively between zebrafish and mammalians, and CRISPR-Cas9 RNP targeting resulted in a similar loss of function as homozygous mutation and MO KD in zebrafish larvae.

Despite the autophagy deficiency was identified in zebrafish with various *atgs* mutations, our work showed that the hematopoietic alterations varied between *atgs*. The findings implied that their regulator effects are, at least partially, canonical autophagy-independent as observed in previous mice studies [9–13]. However, little is known about the specific core ATG-dependent hematopoietic effects in mice since none of them included a comparison between ATGs, and *becn1* and *atg5* are the only two core *atgs* out of six that were selected by this study that have been reported in previous mice studies. The present study determined that *becn1* mutation induced the expansion of HSCs during zebrafish embryonic development to larvae and transiently increased leukocytes or neutrophils. While the up-regulator effects of *becn1* KD on HSCs has been reported in zebrafish with *kri1l* mutation [31], our work for the first time identified the sole role of zebrafish *becn1* in the expansion of HSCs in normal hematopoiesis, which can be ascribed to the directly regulated proteins of HSCs maintenance and disturbances among autophagy, apoptosis, and differentiation [32]. For instance, *SWI/SNF related, matrix associated, actin dependent regulator of chromatin, subfamily c, member (smarcc) 1a* orthologous to human SMARCC1, which was previously shown as a positive regulator for zebrafish HSCs [33], was up-regulated following *becn1* mutation. However, the level of HSCs, including long term-HSC and HSC-containing Lin-Sca-1+c-Kit+ cell, elevated in the spleen while declined in bone marrow in adult mice with *Becn1* CKO in hematopoietic cells [12]. Mouse HSCs reside in both spleen and bone marrow, whereas the HSCs largely resided in the CHT in zebrafish larvae. Therefore, whether Becn1/*becn1* behaves heterogeneously in zebrafish and mice and the precise roles of *becn1* mutation requires the study on HSCs in the spleen and kidney marrow of adult zebrafish. Besides, this discrepancy between mice and zebrafish with *Becn1/becn1* mutations can also be attributed to the unaltered autophagy and autophagic flux in HSCs of mice with *Becn1* CKO [12] and different developmental stages. Strikingly, myeloid cells-specific CKO of Becn1 increased the neutrophil and leukocytes but not the macrophages in mice, which was consistent with our findings in zebrafish, though the autophagic change in myeloid cells was not indicated [34]. Also, a similar expansion of hematopoietic lineages were reported in a fly model with *atg6/becn1* KO [35]. These findings indicated that the expansion of neutrophils and leukocytes in zebrafish with *becn1* mutation may be due to the loss of *becn1* in myeloid cells without affecting the HSCs. In addition to *becn1* mutation, *atg9a* mutation also caused HSCs expansion in zebrafish larvae. Although the study on ATG9a mutation in mammalian HSCs has not yet been reported, a similar proteomic response, especially in HSCs regulation and apoptosis between *atg9a* and *becn1* mutation, was presented in our study, which demonstrated a potential shared mechanism behind the *atg9a* and *becn1* in zebrafish HSCs regulation.

On the other hand, *atg5* mutation showed subtle effects on definitive hematopoiesis, HSCs in particular, in zebrafish larvae and also in mice model with ATG5 or ATG12 (core ATGs of ATG12 conjugation system) CKO, who also harbors a relatively healthy blood system [13]. Nevertheless, hematopoietic abnormalities were observed in mice with ATG5 CKO using a different hematopoietic promoter [9]. The dissimilarity between two mice studies could be explained by the vav:Cre CKO system (with abnormal blood system), which spontaneously effected much earlier than the Mx1:Cre CKO system, which needs the injection of ‘triggers’, such as PolyI:C in an older stage. Moreover, complete loss of *atg5* induced 30% reduction of autophagosomes in the whole zebrafish embryo, whereas it led to an absence of autophagosomes in mouse neonates [36]. Taken together, *atg5* plays a less important role in zebrafish larvae compared with mice; thus *atg5* mutation was associated with a milder hematopoietic phenotype, which was also supported by the trend of change in the number of leukocytes and autophagic vacuoles in leukocytes with *atg5* mutation. In addition, mutation of *atg3* that belongs to another conjugation system showed down-regulator effects on leukocytes, including both neutrophils and macrophages, which is in accordance with hematopoietic defects observed in ATG7-CKO mice probably due to the involvement of ATG7 in both ATG12 and LC3-PE conjugation systems [10]. Although the HSCs were reduced in adult mice with ATG7 CKO, recent work has revealed that ATG7 CKO has no effect on the number and stemness of neonatal HSCs [14]. Similarly, HSCs remained unchanged in zebrafish larvae with either *atg5* or *atg3* mutation. Furthermore, both *atg5* and *atg3* mutations attenuated the number of myeloid cells; in contrast, an expansion of myeloid cells accompanied with lymphopenia was commonly found in adult *Atg7-CKO* mice [10]. Thus, core *atgs* that belong to the core conjugation systems may act distinctly from the ATG7 in myeloid cells. This dissimilarity could also be ascribed to the varied developmental stages and mutations in different hematopoietic lineages as no adverse phenotype was manifested in mice with myeloid cell-specific ATG7 CKO [37]. Besides, a previous study also reported the cell-autonomous effect of FIP200, which together with ATG13 constitute the core autophagy machinery (ULK1 complex), CKO on mouse hematopoiesis. Loss of fetal HSCs and myeloid expansion were characterized in mice with FIP200 CKO [11]. Comparably, we found the *atg13* mutation also elicited transiently decreased HSCs and increased leukocytes, as well as the skewing from macrophages to neutrophils. In summary, mutation of ATGs that belongs to the same autophagy machinery shared the feature of hematopoietic abnormalities, though more evidence is still needed.

In addition to the distinct effects of *atgs* on HSCs, leukocytes, and myeloid cells (neutrophils and macrophages), erythrocytes and myeloid progenitor cells showed similar responses to various *atg* mutations, which may be regulated through the canonical autophagy pathway. Specifically, myeloid progenitor cells were down-regulated, while erythrocytes remained constant. However, their responses in mice differed between ATGs CKO [9–13], which is potentially due to developmental stage difference [14]. More importantly, we determined the interactions between *atgs*, such as the synergistic effect of *atg13+becn1*, *atg9a+atg2a*, and *atg9a+atg3* double mutations on HSCs expansion, which demonstrated the cooperation rather than sloe action of some of the ATGs in definitive hematopoiesis; and highlighted the essentiality of study with multiple ATGs.

However, only one study with double ATGs KO was documented in mice, which revealed an indispensable Ulk1-mediated Atg5-independent autophagy in the regulation of erythropoiesis [38], while the interactions between other ATGs in other lineage hematopoiesis remain concealed. Despite our work providing a more comprehensive picture of various core ATGs in the regulation of definitive hematopoiesis than previous studies, the limitations of this study cannot be neglected: 1) the impact of single *atg* mutation is limited on zebrafish definitive hematopoiesis; 2) cell-autonomous and non-cell-autonomous effects of *atgs* cannot be distinguished, 3) the effect of *atgs* on adult hematopoiesis remains unclear, and 4) more ATGs and combinations are needed, which ultimately calls for the use of inducible hematopoietic cell-specific multiplex *atgs* KO in future studies.

## Materials and methods

### Zebrafish strains and husbandry

Transgenic and wild-type zebrafish were maintained in 14:10 h light:dark cycle and fed brine shrimp twice daily. Tg(Lc3:GFP) [39], Tg(coro1a:DsRed) [40], Tg(cmyb:GFP), Tg(mpx:GFP), and Tg(mfap4:turquoise2) zebrafish lines were used in this study. Tg(mfap4:turquoise2) zebrafish line was generated by micro-injection of pDEST mfap4:turquoise2 plasmid construct (Addgene, 135218; David Tobin Lab) with *in vitro* transcribed tol2 transposase mRNA (Addgene, 31831; Stephen Ekker Lab). Zebrafish embryos were raised at 28.5°C and staged by day post□fertilization (dpf) and morphological criteria as previously described [41]. All animal experiments were performed in accordance with protocols approved by Animal Subjects Ethics Sub-Committee (ASESC) of The Hong Kong Polytechnic University.

### CRISPR-Cas9 sgRNP complex targeting

Target sites of zebrafish *atgs* (*atg13, becn1, atg9a, atg2a, atg5*, and *atg3*) and sgRNAs were identified and designed using Alt-R^®^ CRISPR-Cas9 guide RNA design tool (Integrated DNA Technologies) and CRISPRscan [42]. Target sites were chosen that 1) a high on-target score, 2) a low off-target score, and 3) a restriction enzyme site around the protospacer adjacent motif (PAM) sequence were reported. Target sequence and restriction enzyme site of *atgs* sgRNA used in this study were listed in **Figure 1C and Figure S1E**. Given the target sequences of sgRNAs, CRISPR-Cas9 ‘single guide RNPs’ (sgRNPs) was generated and delivered to zebrafish embryos as previously described [18]. Briefly, synthetic Alt-R^®^ CRISPR-Cas9 sgRNA (Integrated DNA Technologies) was folded at 95°C for 5 min and then cooled down to the room temperature. Alt-R^®^ S.p. Cas9 Nuclease V3 (Integrated DNA Technologies, 1081059) diluted in Cas9 working buffer (20 mM HEPES, 150 mM KCl, pH 7.5) and subsequently was assembled with folded sgRNA in 37°C for 10 min. Around 2nl sgRNP complex was delivered into the cell of zebrafish embryos at one-cell stage through microinjection. For co-injection, two sgRNPs was formed individually and then mixed before micro-injection. Around 4nl sgRNP complexes were injected into the cell of zebrafish embryos. Almost no toxic effect of these sgRNP complexes was observed in zebrafish embryos after microinjection.

### Detection of mutagenesis

Mutagenesis and mutagenic efficiency were detected by using RFLP assay as previously described [43]. Briefly, a 300-600bp PCR fragment covering the designed sgRNA target site was amplified for each *atgs* using corresponding primers listed in **Table S1**. Next, restriction enzyme (listed in **Figure S1E**, New England Biolabs) was then used to cleave the PCR fragment, in which uncleaved band suggested a destroyed restriction enzyme site by sgRNP-induced mutations, while the cleaved band contained wild-type sequence. Mutagenic efficiency was calculated by dividing the intensity of uncleaved band with the intensity of total band measured by ImageJ (NIH). To further confirm the mutations, uncleaved band was sub-cloned into pGEM^®^-T Easy Vector (Promega Corporation, A1360) and the insertions and deletions (indels) were identified by Sanger sequencing.

### Western blotting

The protein level of Lc3 and GAPDH in zebrafish embryos was measured using western blot as previously described [39]. Briefly, total protein extracted from dechorionated and deyolked embryos was resolved on 12% gels (Bio-Rad Laboratories, 1610175) and then transferred to the PVDF membrane (Merck Millipore, IPVH00010). After blocking with 5% nonfat dried milk (Bio-Rad Laboratories, 1706404), membrane was probed with anti-Lc3b (Abcam, ab48394) and anti-GAPDH (Cell Signaling Technology, 2118) primary antibodies. Afterward, the membrane was probed with goat anti-rabbit IgG secondary antibody (Invitrogen, 32460) and visualized using SuperSignal™ West Femto Maximum Sensitivity Substrate (Thermo Fisher, 34095) after wash with with Tris-buffered saline (50 mM Tris base, 150 mM NaCl, pH 7.5) plus Tween-20 (Bio-Rad Laboratories, 1610781).

### LysoTracker Red and Cyto-ID staining

Live zebrafish embryos was incubated with LysoTracker Red DND-99 (Invitrogen, L7528) at 10 μM in E3 medium (5 mM NaCl, 0.17 mM KCL, 0.33 mM CaCl, and 0.33 mM MgSO4, pH 7.4) for 30-45 min at 28.5°C in dark [39]. On the other hand, autophagic vacuoles and nucleus in sorted cells were immediately stained using CYTO-ID^®^ Autophagy detection kit 2.0 with Hoechst 33342 Nuclear Stain (Enzo Life Sciences, ENZ-KIT175) at 37°C in dark following the manufacturer’s instructions [19]. Wash with E3 medium and 1X assay buffer were applied to both LysoTracker Red and Cyto-ID live staining, respectively, before fluorescent microscopic imaging.

### Fluorescence-activated cell sorting

Single-cell suspension for FACS was prepared as described previously [44]. Tg(coro1a:DsRed) zebrafish embryos at 3 dpf were digested with Gibco™ Trypsin-EDTA (0.05%) (Thermo Fisher, 25300062) for 15 minutes at 28°C and then disassociated with pipetting on ice. After termination of Trypsin with CaCl2 (2mM), the suspension was filtered using 40 μm cell strainer (BD Biosciences, 352340) and washed with phosphate-buffered saline (PBS) (VWR Life Science, E404-200TABS) with 1% (vol/vol) fetal bovine serum (FBS) (Thermo Fisher, 26140079). FACS of *coro1a:DsRed+* leukocytes was then conducted in BD FACSAria III Cell Sorter according to the manufacturer’s instructions. Around 8,000 *coro1a:DsRed+* cells was targeted to be sorted using a purity sort mode.

### Fluorescent microscope imaging

Fluorescent images of transgenic zebrafish embryos with or without LysoTracker Red staining or sorted *coro1a+* cells stained with Cyto-ID and Hoechst were taken by using Zeiss Lightsheet Z.1 Selective Plane Illumination Microscope, Leica TCS SPE Confocal Microscope, or Nikon Stereomicroscope with a Nikon DS-Fi2 Camera as previously described [39]. Zebrafish embryos were mounted in 1.5% low-melting agarose (Sigma-Aldrich, A9045) into 35 mm glass-bottom confocal dish or glass capillary before imaging. Tricaine (Sigma-Aldrich, A5040) at 0.16 mg/ml in E3 medium was used as anaesthetic for zebrafish embryos. In addition, Tg(Lc3:GFP) zebrafish embryo was treated with chloroquine (Selleckchem, S4157) at 100 μM in E3 medium before imaging.

### Whole-mount in situ hybridization

Whole-mount in situ hybridization (WISH) was performed on zebrafish embryos following the standard protocol described previously [44]. DIG-labeled anti-sense probes (*cmyb, spi1, lcp1*, and *hbae1*) were made from the pGEM^®^-T Easy vector (Promega Corporation, A1360) containing the gene-coding sequences via *in vitro* transcription using DIG RNA Labeling Kit (Roche, 11175025910). A short bleaching was used to remove the pigments from the fixed zebrafish embryos.

### Mass spectrometry-based proteomics

Total protein was extracted from thirty zebrafish embryos using cell lysis buffer (Sigma-Aldrich, C3228). Purified protein was then digested into peptides using Trypsin (Promega, V5111) and desalted using Pierce C18 Spin Columns (Thermo Fisher, 89870). A label-free quantitative proteomics was conducted on Thermo Fisher Orbitrap Fusion Lumos Mass Spectrometer coupled with Dionex UltiMate 3000 RSLCnano. Identification and quantification was processed with Progenesis QI software, and the abundance of proteins was quantified based on three independent experiments and normalized based on total protein. Arbitrary fold change cut-offs of > 1.5 and significance p-values of 0.05 were set for significantly up-regulated or down-regulated proteins.

### Quantification and statistics

The number of Lc3+ puncta in the muscle was counted in Zeiss ZEN software following the criteria and protocol described in our previous study [39]. Cyto-ID+ autophagic vacuoles was defined by vacuole-like GFP+ fluorescence signals that were distinguished from the background and a similar size of autophagic vacuoles was considered in all the autophagic vacuoles counting in the cells. In addition, ImageJ (NIH) was utilized to measure the relative intensity of proteins in western blot and straighten the WISH images. Data are reported as mean ± standard deviation (S.D.). One-way ANOVA, two-way ANOVA, *C^2^* test, and independent t-test were performed where appropriate using Statistical Package for the Social Sciences (SPSS) Version 14.0 and a *p*-value less than 0.05 was considered statistically significant.

## Abbreviations

ATGs: autophagy-related genes
BECN1: Beclin1
CHT: caudal hematopoietic tissue
CKO: conditional knockout
CQ: Chloroquine
CRISPR: Clustered Regularly Interspaced Short Palindromic Repeats
dpf: days post fertilization
FIP 200: FAK family-interacting protein of 200 kDa
HSCs: hematopoietic stem cells
KD: knockdown
LC3: microtubule-associated protein 1A/1B-light chain 3
MO: Morpholino
PI3K: class 3 phosphoinositide 3-kinase
RFLP: restriction fragment length polymorphism
RNP: ribonucleoprotein
ULK: unc-51 like autophagy activating kinase
WISH: whole-mount in suit hybridization

## Acknowledgements

The zebrafish maintenance was supported by Fish Model Translational Research Laboratory (HTI, PolyU). Light-sheet and confocal fluorescent microscopic imaging, flow cytometry, and mass spectrometry-based proteomics were supported by University Research Facility in Life Sciences (ULS, PolyU).

